# mosaicFlye: Resolving long mosaic repeats using long error-prone reads

**DOI:** 10.1101/2020.01.15.908285

**Authors:** Anton Bankevich, Pavel Pevzner

## Abstract

Long-read technologies revolutionized genome assembly and enabled resolution of *bridged repeats* (i.e., repeats that are spanned by some reads) in various genomes. However, the problem of resolving *unbridged repeats* (such as long segmental duplications in the human genome) remains largely unsolved, making it a major obstacle towards achieving the goal of complete genome assemblies. Moreover, the challenge of resolving unbridged repeats is not limited to eukaryotic genomes but also impairs assemblies of bacterial genomes and metagenomes. We describe the mosaicFlye algorithm for resolving complex unbridged repeats based on differences between various repeat copies and show how it improves assemblies of the human genome as well as bacterial genomes and metagenomes. In particular, we show that mosaicFlye results in a complete assembly of both arms of the human chromosome 6.

## Introduction

Single-Molecule Sequencing technologies (such as the ones developed by Pacific Biosciences (PB) and Oxford Nanopore Technologies (ONT)) opened a new long-read era in genome assembly. Long reads often bridge long repeats and resolve many segmental duplications that otherwise were nearly impossible to assemble using short-read technologies. As a result, the contiguity of the recently described long-read human genome assemblies already exceeds the contiguity of the reference human genome assembled using short reads (Jain et. al, 2018, Miga et al., 2019).

Although long reads greatly improved the contiguity of genome assemblies, resolving long repeats remains a challenging task. For example, the state-of-the-art long-read assemblers fail to fully assemble ∼50% of bacterial genomes from the NCTC 3000 project aimed at sequencing 3000 bacterial genomes from England’s National Collection of Type Cultures (Kamath et al., 2017, Marijon et al, 2019). Additionally, the base-pair accuracy of the long-read assemblies in the repeated regions is reduced (as compared to unique regions) since it is often unclear how to align reads to various repeat copies even if the repeat itself was bridged by some but not all reads.

Long error-prone reads and short accurate reads have their strengths and weaknesses with respect to repeat resolution, e.g., short reads may resolve some repeats that are difficult to resolve with long reads. For example, diverged copies of a long repeat (e.g., copies differing by 3%) often don’t share *k*-mers (for typical *k*-mer size used in short-read assemblers) and thus are automatically resolved by the *de Bruijn graph*-based assemblers such as SPAdes (Bankevich et al., 2012). In contrast, long-read assemblers face difficulties resolving such repeats since repeat copies with a 3% divergence are difficult to distinguish using the error-prone reads that have error rates exceeding 10%. Thus, long-read assemblers trade the ability to resolve the unbridged but divergent repeat copies for the ability to resolve bridged repeats.

Since nearly all genomes have long repeats, long-read assemblers (such as Falcon (Chin et al., 2016), Canu (Koren et al., 2017), Marvel (Nowoshilow et al., 2018), Flye (Kolmogorov et al., 2019a), wtdbg2 (Ruan and Li, 2019), and others) currently face the same repeat-resolution challenge that short-read assemblers faced a decade ago, albeit at a different scale of repeat lengths. To improve the contiguity of assemblies, long read technologies are often complemented by Hi-C (Ghurye et al., 2017) and optical mapping (Weissensteiner et al., 2017) data. However, these technologies add significant cost to the sequencing projects (estimated to be higher than the cost of generating long reads in a typical vertebrate assembly project) and typically fail to accurately reconstruct various repeat copies even if they resolve these copies. Moreover, it remains unclear how the inherent errors of data generated by these additional technologies affect the accuracy of the final assemblies. Thus, resolving unbridged repeats using long reads represents the crucial step towards improving the long-read assembly algorithms (Bongartz et al., 2018) and achieving the goals of large sequencing programs such as the Earth BioGenome Project (Lewin et al., 2018).

Repeats in a genome accumulate mutations and result in divergent repeat copies, e.g., most segmental duplications in the human genome diverge by more than 1% (Pu et al., 2018). Vollger et al., 2019 described how to use the variations between various repeat copies for reconstructing all copies of a divergent repeat. This problem is similar to the haplotype assembly problem in the case of high ploidy with several important distinctions. First, the number of copies (edge multiplicity in the assembly graph) is often unknown and may vary along a repeat. Second, unlike the haplotype assembly, where all haplomes align to a consensus sequence, many repeats have complex mosaic structure (Pevzner et al., 2004, Jiang et al., 2007, Pu et al., 2018) that prevents utilization of a single consensus sequence as a template for aligning all copies of a repeat. Such mosaic repeats are also common in cancer genomes, making it difficult to analyze duplications that represent the hallmarks of many cancers (Nattestad, 2018). Such difficult cases were not considered in Vollger et al. 2019 that focused on resolving repeat copies with a *single* consensus sequence. However, *mosaic repeats* consist of several smaller sub-repeats that appear with varying multiplicities and in different combinations within various copies of a mosaic repeat. Such mosaic repeats are common, e.g., most segmental duplications in the human genome (Pu et al., 2018) and many repeats in bacterial genomes (Pevzner et al., 2004) represent mosaic repeats.

We describe mosaicFlye algorithm for reconstructing individual copies of mosaic repeats based on differences between various copies and demonstrate how it contributes to improving genome and metagenome assemblies. The mosaicFlye code is available at https://antonbankevich.github.io/mosaic/.

## Methods

### De Bruijn graph

Given a parameter *k*, we define the *genome graph* by representing each chromosome of length *n* in the genome as a path on *n-k+*1 vertices (a position in the chromosome corresponds to a vertex labeled by a *k*-mer that starts at this position). Let *DB*(*Genome, k*) be the de Bruijn graph of a genome *Genome*, where vertices and edges correspond to *k*-mers and (*k+*1)-mers in *Genome*, respectively (Compeau et al., 2011). Alternatively, the de Bruijn graph can be constructed by “gluing” identical *k*-mers in the genome graph (Pevzner et al., 2004). We will work with the *condensed de Bruijn graphs*, where each *non-branching* path is collapsed into a single edge labeled by the corresponding substring of the genome. Each chromosome in *Genome* corresponds to a path in this graph and the set of these paths forms the *genome traversal* of the graph.

Given a read-set *Reads* sampled from *Genome*, one can view each read as a “mini-chromosome” and construct the de Bruijn graph of the resulting genome (Pevzner et al., 2004) that we refer to as *DB*(*Reads, k*). In contrast to *DB*(*Genome, k*) (where each edge is labeled by a substring of *Genome*), edges of *DB*(*Reads, k*) inherit errors in reads. Since the graph *DB*(*Reads, k*) encodes all errors in reads, it is much more complex than the graph *DB*(*Genome, k*). In the case of short reads, various graph-based error correction approaches transform the graph *DB*(*Reads, k*) into the *assembly graph* that approximates *DB(Genome, k)* (Pevzner et al., 2004, Bankevich et al., 2012). However, these error correction approaches assume that nearly all *k*-mers from *Genome* also occur in reads, the condition that holds for short-read datasets but is violated for long reads. As a result, constructing an accurate assembly graph from long error-prone reads is a challenging problem.

### Repeat graph

Kolmogorov et al., 2019a developed a Flye assembler that attempts to solve this problem by making some concessions. Flye constructs the *repeat graph* of long reads (also known as the *A-Bruijn graph*) with the goal to approximate the de Bruijn graph *DB*(*Genome, k*) in the case of a large *k*, e.g., *k*=1500. Since this task proved to be difficult in the case of error-prone reads, the Flye assembler collapses similar (rather than only identical as in the de Bruijn graph) *k*-mers in the genome graph into a single vertex in the repeat graph and labels this vertex by the consensus sequence of all collapsed *k*-mers.

Specifically, to construct the repeat graph of a genome, Flye generates all local self-alignments of the genome against itself that have divergence below the *divergence threshold d*%. Two positions in the genome are defined as *equivalent* if they are aligned against each other in one of these alignments. Pevzner et al, 2004 defined the *repeat graph RG*(*Genome, k, d*) as the graph obtained from the genome graph by collapsing all equivalent positions (vertices) into a single vertex (Kolmogorov et al., 2019a used a similar construction of the repeat graph). Note that the graph *RG*(*Genome, k, 0*) = *DB*(*Genome, k*).

Kolmogorov et al., 2019 defined the repeat graph *RG*(*Reads, k, d*) similarly to *RG*(*Genome, k, d*) by applying the same approach to a “genome” formed by all reads (each read is viewed as a “mini-chromosome”). They further described how to construct *RG*(*Reads, k, d*) in the case when *d* is not too small (e.g., exceeds 5%) and demonstrated that *RG*(*Reads, k, d*) approximates *RG*(*Genome, k, d*). However, although the problem of constructing (approximating) the repeat graph of a genome from long error-prone reads has been solved, it remains unclear how to construct the de Bruijn graph of a genome from such reads. Solving this problem is arguably one of the most pressing needs in the assembly of long error-prone reads since it would result in assemblies of the same quality as assemblies of long *error-free* reads.

Figure 1 illustrates this problem in the case of a genome where each *k*-mer is unique. In this case, the de Bruijn graph is a cycle (resulting in a unique genome reconstruction) but the repeat graph is more complex since the genome contains similar *k*-mers.

**Figure 1.**
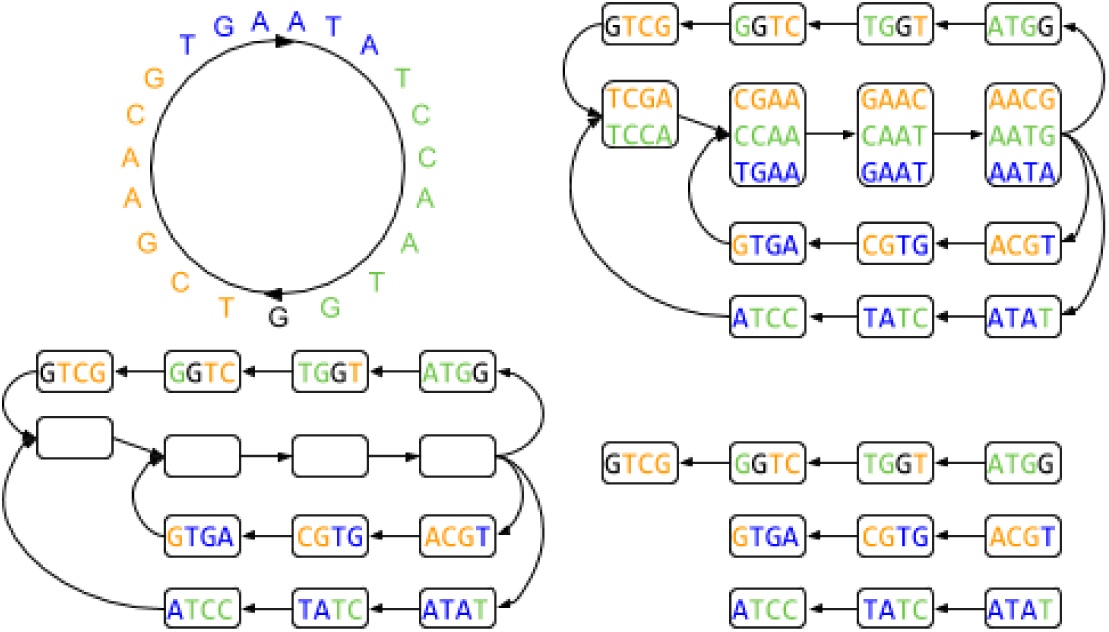
A genome (top left), its repeat graph (top right), its cryptic repeat graph (bottom left), and its partial repeat graph (bottom right). Since all 4-mers in the genome are unique, its de Bruijn graph is a cycle. However, its repeat graph is not a cycle (two 4-mers are defined as similar if they are at most 1 substitution apart). In the realm of genome assembly (when the genome is unknown), the label of each complex vertex in the cryptic repeat graph represents the consensus of all individual *k*-mers that were glued into this vertex. Since this consensus *k*-mers do not reveal information about the individual k-mers, we assume that the labels of complex vertices in the cryptic repeat graph are unknown. The partial repeat graph (a subgraph of the repeat graph formed by its simple vertices), provides even less information than the cryptic repeat graph.

mosaicFlye uses variations between various copies of a mosaic repeat for resolving these copies and thus untangling the repeat graph of reads *RG*(*Reads,k,d)* constructed by the Flye assembler. Kolmogorov et al., 2019a first described how Flye constructs the graph *RG*(*Genome,k,d*) and later explained how to construct the graph *RG*(*Reads,k,d*) by simply applying the same approach to a “genome” formed by all reads. Similarly, we first describe the idea of the mosaicFlye algorithm using *Genome* and later explain how it works in the case when *Genome* is unknown and only the read-set *Reads* is given.

### The challenge of transforming the repeat graph into the de Bruijn graph

Although the repeat graph constructed by Flye provides a useful representation of repeats in a genome, its main deficiency (compared to the graph *DB*(*Genome, k*)) is that sequences of various instances of each repeat edge remain unknown as they are substituted by a consensus sequence of this edge. Flye and other long-read assemblers do not resolve such repeats, resulting in a lower contiguity as compared to the assembly represented by the de Bruijn graph *DB*(*Genome, k*).

Figure 2 illustrates the differences between the de Bruijn graph and the repeat graph in the case of a “genome” that contains three instances of a mosaic repeat. Colored segments of the genome represent non-diverged parts of the three copies of a mosaic repeat (each triple of colored segments is collapsed in a single colored edge in the de Bruijn graph). In the repeat graph, all three instances of the mosaic repeat are collapsed into a single purple edge, resulting in a simpler graph (as compared to the de Bruijn graph) but making it difficult to reconstruct three different instances of the mosaic repeat.

**Figure 2:**
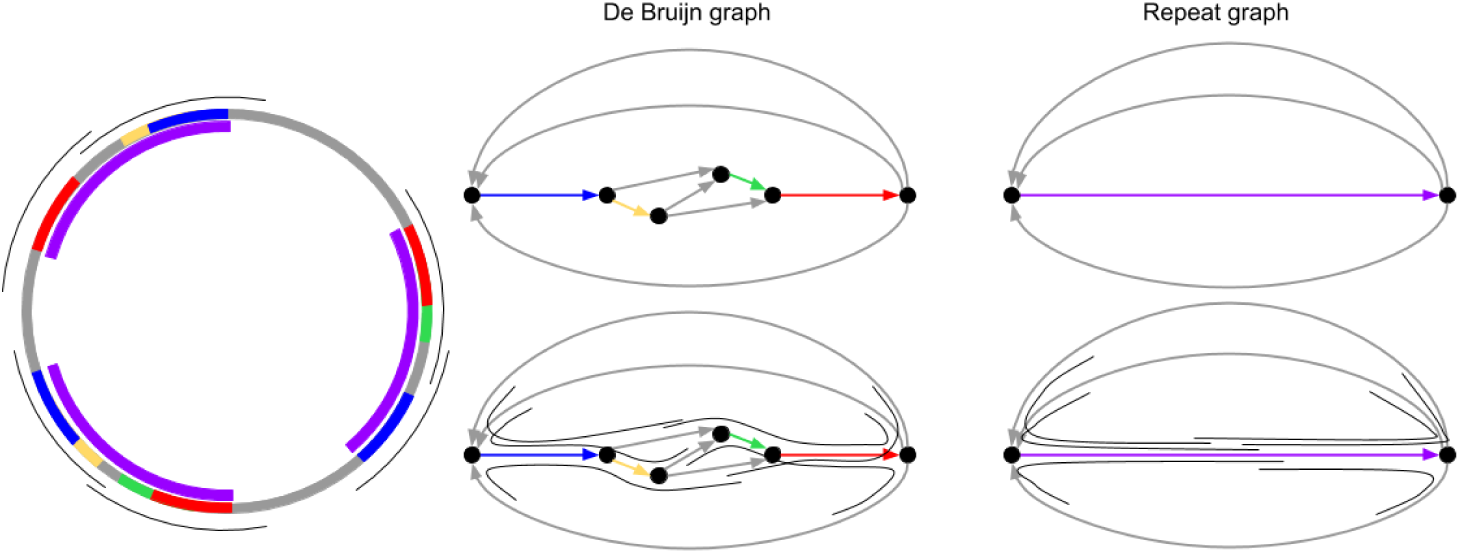
A genome with three copies of a mosaic repeat (left), its de Bruijn graph (middle top), its repeat graph (right top), alignment of reads to the de Bruijn graph (middle bottom), and to the repeat graph (right bottom), The genome contains three instances of a mosaic repeat (marked by purple) formed by four non-diverged sub-repeats shown in red, yellow, green, and blue with 3, 2, 2, and 3 copies, respectively. Six unique regions in the genome are shown in gray. Reads bridging red, yellow, green, and blue sub-repeats are shown in gray.

Flye classifies edges in the repeat graph *RG*(*Reads, k, d*) into *unique* and *repeat* edges (Kolmogorov et al., 2019). A read *bridges* a repeat if its *read-path* in the repeat graph starts at a unique edge, traverses some repeat edges, and ends in a unique edge. Although the repeat-bridging reads enable the *repeat resolution* algorithm in the Flye assembler, this algorithm has some limitations. Figure 2 illustrates that, in contrast to the de Bruijn graph that has bridging reads (that enable complete genome reconstructions), the repeat graph does not have bridging reads since the purple repeat is longer than any read. mosaicFlye addresses this limitation of the repeat graph and enables resolves unbridged repeats as long as they have some diverged positions.

### Genome polishing challenge

Each read *Read* originates from a region in a genome that we refer to as its *origin*. A read is *correctly aligned* to a genome if it is aligned against its origin and *incorrectly aligned* otherwise. A read-set is called *correctly aligned* if all reads in this set are correctly aligned. We define a *high-coverage* read-set as a read-set with coverage depth exceeding the *coverage* threshold (the default value is 30x in the case of PB reads).

Even if a genome is unknown and only its error-prone version is given, *polishing* algorithms correct most errors and generate highly accurate genome sequence in the case of high-coverage and correctly aligned read-sets (Loman et al., 2015, Lin et al., 2016, Vaser et al., 2017, Lima et al., 2019). However, it is not clear how to construct a draft genome sequence, not to mention correctly align all reads. Although it is easy to correctly align reads that fit into unique regions of a genome (or overlap these regions), it is not clear what specific copy of a repeat in a draft genome a read correctly aligns to. Incorrect read alignments to “wrong” repeat copies result in contamination of reads recruited to each copy by “foreign” reads from other copies that mislead the polishing procedure and turn multiple diverged copies of a repeat into a single consensus of all copies. On the other hand, if the entire error-free genome sequence is known, finding correct read alignments turns into an easy problem since the alignment with the maximal score is typically correct (unless the read aligns to identical or nearly identical instances of the repeat, which makes selection of the correct alignment impossible). Thus, correct read alignments are required to polish the draft genome but the polished genome is needed to construct the correct read alignments.

### Transforming the repeat graph into the de Bruijn graph by gradually shrinking the set of similar *k*-mers

To explain the idea of the mosaicFlye algorithm, we first describe how it transforms the repeat graph of *Genome* into the de Bruijn graph of *Genome* (we later explain how it works in the case when *Genome* is unknown and only the read-set *Reads* is given). Flye uses a rather general concept of similarity that takes into account mismatches, insertions, and deletions. For the sake of simplicity, we will consider a less general notion of similarity (limited to mismatches only) when similar *k*-mers are defined as *k*-mers with at most *δ* mismatches. We will thus redefine the repeat graph for this less general notion using the concept of the similarity graph.

Given a set of *pairs* of *k*-mers from a genome (referred to as *sim*=*sim*(*Genome*)), we define a *similarity graph* with the vertex-set formed by all *k*-mers in *Genome* and the edge set *sim*(*Genome*). We call two *k*-mers *similar* if they belong to the same connected component of the similarity graph. In difference from the de Bruijn graph (constructed by gluing all *identical k*-mers into a single vertex), the *repeat graph RG*(*Genome*)=*RG*(*Genome,sim*) is constructed by gluing all *similar k*-mers in the genome graph into a single vertex. Note that the de Bruijn graph is the repeat graph with the empty set *sim(Genome)*.

The number of vertices in the repeat graph is equal to the number of connected components in the similarity graph and each vertex *v* is labeled by a set of *k*-mers (forming a connected component in the similarity graph) that we refer to as *label*(*v*). A vertex is called *simple* if its label consists of a single *k*-mer, and *complex* otherwise.

In the realm of long-read assembly, although *Genome* is unknown, the Flye assembler constructs the repeat graph *RG*(*Reads*)*=RG*(*Reads,sim*) from reads that approximates the graph *RG*(*Genome*)*=RG*(*Genome,sim*). For the sake of simplicity, in addition to the condition that all *k*-mers with at most *δ* mismatches form the set *sim*(*Genome*), we also assume that every two *k*-mers in the same connected component of the similarity graph differ by at most *δ* mismatches (*transitivity condition*).

We will transform the repeat graph *RG*(*Reads*) (that approximates *RG*(*Genome*)) into a graph that approximates the de Bruijn graph *DB*(*Genome*)*=DB*(*Genome,k*) by gradually shrinking the set *sim* and thus resolving more and more repeats. We note that, since *Genome* is unknown, we will perform this transformation using some operations that can be implemented using information about reads.

### Ungluing complex vertices in the repeat graph

Each complex vertex *v* in the repeat graph corresponds to a connected component in the similarity graph. Removal of all edges of this connected component results in a smaller set of *k*-mer pairs that we refer to as *sim(v).* Given a complex vertex *v* in the repeat graph *RG*(*Genome,sim*), the *ungluing operation* on this vertex returns the repeat graph *RG*(*Genome,sim*(*v*)). Below we show how to use reads to perform some ungluing operations even though, in the realm of genome assembly, the labels of complex vertices are unknown.

### Cryptic repeat graphs

Flye classifies edges of the graph *RG*(*Reads*) into unique and repeated and further applies the Flye polishing algorithm (Lin et al., 2016) to derive the consensus of each edge. Since polishing results in an accurate consensus in the case of a unique edge, we assume that these edges are perfectly polished and thus all *k*-mers on these edges (referred to as *unique k*-mers) are known and represent *k*-mers from *Genome*. In contrast, since the polishing of repeated edges results in a consensus sequence (rather than sequences of individual instances of each repeat), we assume that the labels of *repeated k-mers* (located on repeat edges) are unknown.

Thus, to model the reality of genome assembly, we will work with *cryptic repeat graphs* with known labels of simple vertices but hidden labels of complex vertices (Figure 1). Our goal is to transform a cryptic repeat graph into the de Bruijn graph, an easy task if we were able to perform a series of ungluing operations on all vertices of the cryptic repeat graph. Although it is not clear how to perform ungluing operations in the realm of genome assembly, mosaicFlye uses reads to identify some vertices in the cryptic repeat graph that enable ungluing operations.

### Transforming a cryptic repeat graph into a de Bruijn graph

A complex vertex is called *semi-complex* if either all its predecessors are simple or all its successors are simple. An ungluing operation on a semi-complex vertex is called a *legal ungluing operation*. mosaicFlye iteratively identifies semi-complex vertices in the repeat graph, performs legal ungluing operation on these vertices, and returns the resulting graph that we refer to as *mosaic*(*Genome*). For now, we treat a legal ungluing operation as a black-box function (without explaining how it works) but the subsection “Implementing legal ungluing” explains how mosaicFlye uses reads to implement this black-box.

Legal ungluing operation contributes to repeats resolution by reducing the number of complex vertices in the repeat graph. However, although the graph *mosaic(Genome)* has no semi-complex vertices (otherwise, we would be able to perform additional ungluing operations) it may still have complex vertices. We refer to the subgraph on the set of remaining complex vertices in *mosaic*(*Genome*) as the *complex subgraph*. Since there are no sources and sinks in the complex subgraph (otherwise, a source or a sink vertex would be semi-complex), each connected component of the complex subgraph contains a directed cycle. Such connected components represent the most complex mosaic repeats (referred to as *cyclorepeats*) such as ultralong tandem repeats or cyclic segmental duplications analyzed in Pu et al., 2018. With the exception of the recently proposed algorithm for assembling centromeres (Bzikadze and Pevzner, 2019), existing repeat resolution tools (including mosaicFlye) are unable to resolve cyclorepeats.

### From *mosaic(Genome)* to *mosaic(Reads)*

Above we explained how to construct the graph *mosaic*(*Genome*) but did not explain what various concepts introduced above (e.g., the “legal ungluing” black-box) mean in the realm of genome assembly. In the next section, we explain how mosaicFlye constructs the graph *mosaic*(*Reads*).

To construct the repeat graph of a genome, Flye generates a *genomic dot-plot* representing all local self-alignments within a genome (analog of the similarity graph) and uses this dot-pot to construct the graph *RG(Genome)* (analog of the graph *RG*(*Genome, sim*)). Similarly, to construct the repeat graph of reads, it generates all local alignments between reads (analog of the similarity graph) and constructs the graph *RG*(*Reads*) (analog of the graph *RG*(*Genome,sim*)) by considering each *disjointig* (Kolmogorov et al., 2019a) as a mini-chromosome. mosaicFlye iteratively performs legal ungluing operations on the graph *RG(Reads)* as described below.

### Implementing legal ungluing: from *k*-mers to *K*-mers

mosaicFlye sets two parameters: the *k-mer-size* (typical value *k*=1500) and a larger *K-mer size* (typical value *K*=2500). The parameters *k* and *K* in mosaicFlye are not unlike parameters *k* and *k*+1 in the de Bruijn graph construction. In the case of the de Bruijn graph, each vertex *a* is labeled by a *k*-mer and outgoing edges from *a* are labeled by (*k+*1)-mers. Each such (*k*+1)-mer reveals information about a vertex *b* that follows *a* (lays in the *immediate* vicinity of a vertex *a*). However, information about this (*k*+1)-mer is not sufficient to unglue the vertex *b* since it lacks information about the outgoing edges from *b*. mosaicFlye takes a step further by considering a larger *K*-mer that follows *a* (for *K* > *k*+1) and thus speeding up the ungluing process.

Let *a* be a unique *k*-mer from *Genome* and *extension_K_*(*a*) be the (unknown) *K*-mer in *Genome* with prefix *a* (for *K > k+*1). We define *similar(a)* as the set of all *k*-mers from *Genome* that are similar to *a* and align all reads to *a* in a hope to find *extension_K_*(*a*). However, in the case of error-prone reads, this alignment will return reads spanning *all k*-mers that are similar to *a* rather than all reads spanning a *single k-m*er *a*, making it impossible to reconstruct *extension_K_*(*a*). However, below we show that the situation is not hopeless if all *k*-mers from *similar*(*a*)*=*{*a_1_, …, a_t_*} represent simple and unique vertices in *RG(Reads*). Indeed, if one aligns a read to each *k*-mer in *similar*(*a*), a highest-scoring alignment among these *t* alignments almost always detects a specific *k*-mer in *similar*(*a*) spanned by this read. This observation results in a partitioning of all reads that align to *a* into *t* clusters such that reads from the *i*-th cluster span the *k*-mer *a_i_.* Thus, given a semi-complex vertex *v* in *RG*(*Reads*), one can consider all its predecessors (or successors) and align reads to each of *t* predecessors, thus resulting in *t* clusters of reads. Performing polishing on each of these clusters reveals the set of *K*-mers {*extension_K_*(*a_1_*)*, …, extension_K_*(*a_t_*)} for *K > k+*1. mosaicFlye uses these *K*-mers to unglue the vertex *v*.

### Partial repeat graphs

Above we assumed that the repeat graph is given (even though the labels of its complex vertices are unknown) and described how to transform it into the de Bruijn graph using legal ungluing operations. However, since Flye may distort the structure of complex mosaic repeats in the repeat graph and miscalculate the multiplicities of its subrepeats, the repeat graph constructed by Flye is not necessarily 100% correct. Below we assume that we are only given the subgraph of the repeat graph formed by its unique edges (referred to as the *partial repeat graph* that is illustrated in Figure 1). We use reads to iteratively transform the partial repeat graph (that is reliably reconstructed by Flye) into the de Bruijn graph of the entire genome.

### Transforming a partial repeat graph into a de Bruijn graph

At each iteration, mosaicFlye finds the best-scoring alignment of each read to the partial repeat graph. Given a *k*-mer (vertex) *a* in a partial repeat graph, the reads aligned to *a* reveal a *K*-mer *extension* (*a*). The *k*-mer that starts in the 2^nd^ position of the *K*-mer *extension_K_*(*a*) (i.e., the *k*-mer that follows *a*) may represent a still unexplored complex vertex of the repeat graph. If this vertex (referred to as *next*(*a*)) is semi-complex, it can be unglued as described above, thus adding new simple vertices to the partial repeat graph. The repeated application of this procedure has the potential to eventually reconstruct the entire de Bruijn graph.

However, this procedure faces the challenge of verifying whether a still unexplored complex vertex *nex*t(*a*) of the repeat graph is semi-complex. Although this test is easy to conduct when the entire repeat graph is given, it requires additional analysis in the case of the partial repeat graph. Specifically, mosaicFlye aligns all reads to *next*(*a*) and uses the aligned reads to find all *k*-mers that *precede next*(*a*). If all these *k*-mers represent simple and unique vertices in the partial repeat graph, we classify the vertex *next*(*a*) as semi-complete and unglue it as described above.

### mosaicFlye meets the realities of genome assembly

The description above leaves many questions about constructing the graph *mosaic*(*Reads*) unanswered, e.g., “How do we infer accurate labels of simple vertices in the repeat graph?”, “How do we align all reads to a given *k*-mer *a* to find *extension_K_*(*a*)”, etc. These questions are answered in Appendices that we briefly summarize below.

Appendix “Fitting and overlap alignments of reads” explains how mosaicFlye aligns reads against an assembly and generates accurate labels of simple vertices in the repeat graph. Although reads originating from a unique region in an assembly are easy to align, it is not clear how to align reads originating from the repeated regions (Li, 2018). mosaicFlye generates the sets *Fitting*(*Read, C*) and *Overlap*(*Read, C*) of all high-scoring fitting and overlap alignments between a read *Read* and a contig-set *C*. This procedure results in generating accurate (polished) sequences of simple vertices.

Appendix “Alignment tournament” explains how mosaicFlye selects the correct alignment of a read out of many alignments in *Fitting*(*Read, C*) and *Overlap*(*Read, C*). Since repeats in a genome accumulate mutations, the correct alignment of a read (to a repeat copy that it originated from) typically has a larger percent identity than the alignment of the same read to an incorrect repeat copy. However, this difference is often small compared to the percent identity between the read and its origin and, in the case of highly similar repeat copies, errors in a read sometimes result in cases when the percent identity of the correct read alignment is lower as compared to the incorrect one. To avoid selecting false alignments, mosaicFlye uses a probabilistic model for comparing alignments described in Lin et al., 2016 and specifies the tournament between different repeat copies “competing” for a given read.

Extending *k*-mers into *K*-mers is a key step of mosaicFlye that requires correct read alignments. Section “Genome polishing challenge” explained that correct read alignments are required to polish the draft genome but the polished genome is needed to construct the correct read alignments. To resolve this catch-22, mosaicFlye uses the *expanding alignment-consensus loop* (*EACL*) and applies it for genome polishing (appendix “Alignment-consensus loop”) as well as for transforming the repeat graph into the de Bruijn graph (Appendices “Expanding alignment-consensus loop” and “Expanding alignment-consensus loop for transforming the repeat graph into the de Bruijn graph”).

Figure 3 illustrates an iterative transformation of the repeat graph into the de Bruijn graph using EACL. In the first iteration, mosaicFlye finds all reads (referred to as *incoming* reads) that align to the three incoming unique edges and “enter” into the purple repeat (Figure 3, upper left). Since the incoming reads extend into the repeat edge, they cover the neighborhoods of the three incoming repeat edges in the genome traversal and thus provide information about three different instances of the repeat edge, at least for their initial segments that are well covered by the incoming reads. Since these segments are covered by correctly aligned reads, they can be accurately polished, thus revealing the sequences of each such segment. mosaicFlye replaces the beginning of the purple edge representing these segments by the part of the de Bruijn graph constructed from these segments. As a result, two repeats (blue and orange) and three unique (grey) edges, that are missing in the initial repeat graph, are revealed (Figure 3, upper right). In the second iteration, mosaicFlye generates new incoming reads by aligning reads to the new unique edges constructed at the previous iteration (Figure 3, middle left) and reveals the remaining structure of de Bruijn graph (Figure 3, middle right). Finally, after one more EACL iteration, it generates accurate consensus sequences of all edges within a mosaic repeat and concludes the transformation of the repeat graph into the de Bruijn graph. The length of each repeat edge in the resulting graph has reduced (compared to the length of the initial purple edge), leading to the emergence of bridging reads and thus a possibility to resolve repeats in the resulting de Bruijn graph.

**Figure 3:**
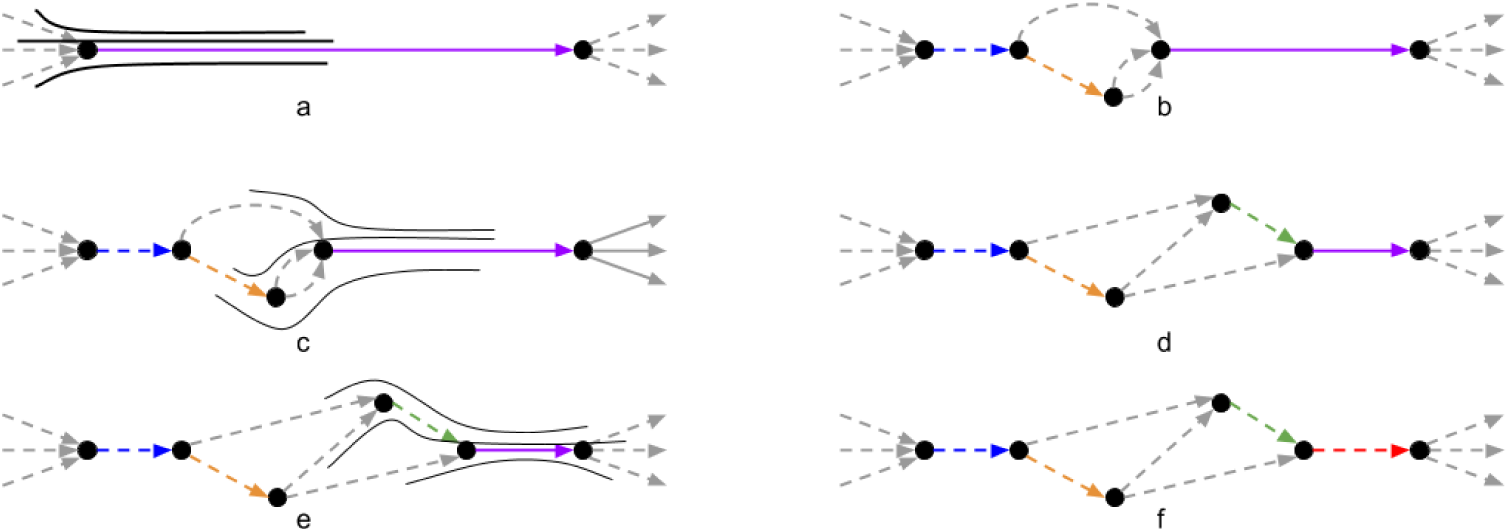
Transforming the repeat graph (top left) into the de Bruijn graph (bottom right). Resolved edges are shown as dashed and unique edges are shown as grey. Paths highlighted by solid lines show the alignment of the consensus of reads that align to unique edges. Note that these paths represent the consensus of reads rather than individual reads as in Figure 2.

The EACL approach faces the challenge of selecting the correct alignment of reads to the growing contigs. Although aligning a read to an initial assembly is an easy task (since all *k*-mers in the initial assembly are unique), it is not clear how to align a read to each intermediate assembly since some *k*-mers in intermediate assemblies are not unique. Indeed, since a read may overlap with several contigs in an intermediate assembly, the choice of the correct overlap alignment becomes non-trivial. Appendix “The challenge of aligning reads to the growing assembly” explains how mosaicFlye addresses this challenge.

Flye reconstructs the accurate sequences of unique edges that mosaicFlye uses as the initial contigs in the EACL approach. It initializes the set of *resolved k-mers* as the set of all *k*-mers occurring in unique edges and iteratively expands it with each EACL iteration in an attempt to find as many resolved *k*-mers as possible. Appendix “Expanding the set of resolved *k*-mers by traversing mosaic repeats” describes how mosaicFlye expands the set of resolved *k*-mers.

mosaicFlye faces difficulties in the case of corrupted mosaic repeats that inaccurately represent various subrepeats of a mosaic repeat. Appendices: “Corrupted mosaic repeats”, “From a resolved prefix *k*-mer to a resolved *K*-mer” and “From a resolved *K*-mer to a resolved suffix *k*-mer” explain how mosaicFlye addresses this challenge. Appendix “Merging contigs and bridging repeats” describes how mosaicFlye merges contigs and bridges repeats.

## Results

### Benchmarking mosaicFlye

Long-read genome assemblers often generate highly contiguous assemblies (as compared to short-read assemblies) that consists of a few contigs. Since mosaicFlye aims to resolve the remaining few unbridged repeats and accurately reconstruct the sequence of each repeat copy, the traditional genome assembly metrics (such as N50 or assembly size) sometimes do not adequately reflect the improvements in the mosaicFlye assembly as compared to the Flye assembly. Thus, the key advantage of mosaicFlye over existing assembly tools lies in reconstructing the accurate sequences of repeat copies and a relatively small reduction in the number of contigs. However, these improvements are important, for example, long bacterial repeats often correspond to important sequences such as 16s rRNAs (Yuan et al., Bioinformatics 2015) or antibiotic resistance genes (Antipov et al., 2019). To benchmark mosaicFlye, we applied it to multiple datasets with complex repeats that the state-of-the-art long-read genome assemblers failed to resolve.

Wick and Holt, 2019 recently benchmarked various long-read assemblers and demonstrated that Flye improves on other long-read assemblers in the case of bacterial genomes. Since this conclusion was further confirmed in recent bacterial studies (Ring et al., 2018, Schmid et al., 2018, Somerville et al., 2019), we only analyzed how mosaicFlye improves on the Flye and metaFlye (Kolmogorov et al., 2019b) assemblies in the case of bacterial genomes and metagenomes. In the case of the human genome, we analyzed both Canu and Flye assemblies.

### Datasets

We benchmarked mosaicFlye on isolate bacterial datasets using datasets from the NCTC 3000 project. We randomly selected 20 datasets that were not assembled into a single circular chromosome by Flye. We also benchmarked mosaicFlye on metagenomic datasets using the ZymoBIOMICS Microbial Community Standards dataset generated using ONT reads (Nicholls et al., 2019). The ZymoEven mock community consists of eight bacteria with an abundance ≈12% and two yeast species with an abundance ≈2%. ZymoBIOMICS community was sequenced using GridION (total read lengths 14 Gb) and PromethION (total read lengths 146 Gb).

Additionally, we analyzed the human genome dataset of ONT reads generated by the T2T consortium (Miga et al., 2019) and released on March 2, 2019. The dataset contains 11,069,717 reads (155 Gb total length, 50x coverage, N50=70 kb) generated from the CHM13hTERT female haploid cell line.

### Resolving repeats in bacterial isolates with mosaicFlye

Table 1 illustrates that mosaicFlye improved assembly of 14 out of the 20 bacterial datasets. Note that some repeats that are not resolved by mosaicFlye have a very low divergence and thus cannot be resolved even in theory. Figure 4 presents examples of repeats resolved by mosaicFlye.

**Figure 4.**
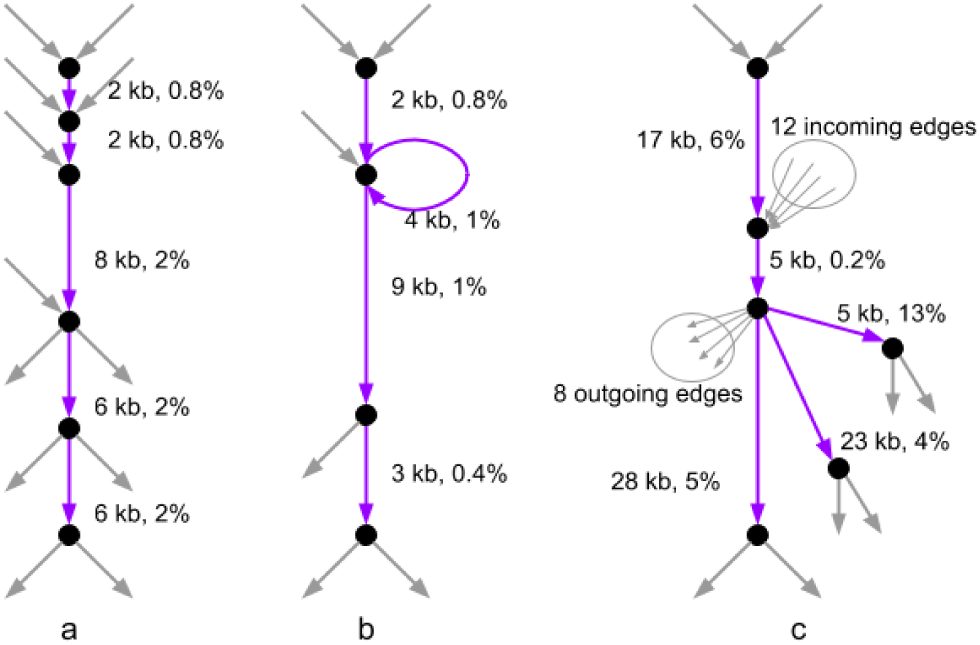
Mosaic repeats in the assembly graphs for the NCTC dataset 10864 (a), the NCTC dataset 9012 (b) and the ZymoEven GridION dataset (c) that were resolved by mosaicFlye. Purple edges represent repeat edges that form mosaic repeats. The unique edges that are adjacent to unique edges are shown in grey. Each repeat edge has two labels showing its length and divergence. The lengths of the longest copies of a mosaic repeat is 25 kb for the NCTC dataset 10864 and 18 kb for the NCTC dataset 9012. The multiplicity of different subrepeats within these repeats varies from 2 to 5 and the divergence between repeat copies varies between 0.4% and 2%. The mosaic repeat in the ZymoEven GridION dataset consists of five subrepeats that are incident to 28 unique regions (shown as black edges). Five subrepeats have lengths 28 kb, 5 kb, 17 kb, 5 kb, and 23 kb.

**Table 1.** Results of mosaicFlye on 20 selected datasets from the NCTC 3000 project. The column “# resolved repeat copies” shows the number of pairs of unique edges from the initial repeat graph that were connected into a single contig by mosaicFlye. This value roughly corresponds to the reduction in the number of contigs in the assembly, with the only difference that a contigs that becomes circular as the result of the mosaicFlye repeat resolution contributes one additional point to this score.

| dataset | # contigs in the Flye assembly | # contigs in the mosaicFlye assembly | # resolved repeat copies | total length of all repeat copies resolved by mosaicFlye (kb) |
| --- | --- | --- | --- | --- |
| 4450 | 35 | 29 | 8 | 66 |
| 7921 | 8 | 5 | 3 | 12 |
| 8294 | 6 | 4 | 2 | 60 |
| 8333 | 18 | 8 | 10 | 136 |
| 8359 | 16 | 8 | 8 | 54 |
| 8603 | 6 | 4 | 2 | 43 |
| 8684 | 10 | 10 | 0 | 0 |
| 9002 | 3 | 3 | 2 | 96 |
| 9006 | 8 | 3 | 5 | 71 |
| 9007 | 4 | 4 | 0 | 0 |
| 9012 | 6 | 4 | 2 | 40 |
| 9016 | 8 | 8 | 0 | 0 |
| 9024 | 4 | 4 | 0 | 0 |
| 9028 | 15 | 10 | 5 | 51 |
| 9657 | 9 | 7 | 4 | 53 |
| 10864 | 14 | 9 | 5 | 73 |
| 11658 | 6 | 6 | 0 | 0 |
| 11692 | 10 | 4 | 6 | 20 |
| 11962 | 5 | 3 | 2 | 15 |
| 12158 | 5 | 5 | 0 | 0 |

### Resolving repeats in metagenomes with mosaicFlye

Since the GridION dataset has a lower coverage than the PromethION dataset, the metaFlye assembly graph of the ZymoEven GridION dataset is more complex than the assembly graph of the ZymoEven PromethION dataset. To challenge mosaicFlye, we applied it to a complex unbridged repeat in the assembly graph of the ZymoEven GridION dataset. Since this repeat was bridged by some reads in the PromethION dataset, we used these bridging reads for checking the accuracy of the mosaicFlye repeat resolution. Using these PromethION reads we verified that mosaicFlye correctly resolved this repeat (Figure 4c) using only reads from the GridION dataset.

### Resolving repeats in the human genome with mosaicFlye

Recently, the Telomere-To-Telomere (T2T) consortium initiated an effort to generate a complete *de novo* assembly of the human genome from ultralong ONT reads (Miga et al., 2019). These reads were assembled by Canu and Flye, resulting in two very similar assemblies with higher contiguity than the hg38 reference human genome. Since no chromosome was assembled into a single contig, the T2T consortium now works on finishing these assemblies (Miga et al., 2019, Bzikadze and Pevzner, 2019).

We selected one of the chromosomes (chromosome 6) that is close to being completely assembled in a fully automated fashion. It has only two unassembled regions: the centromere and a 300 kb long segmental duplication (SD). We applied mosaicFlye to the 300 kb long SD with two copies located at positions 57967000 and 60718600 in the hg38 human genome assembly. Since these two copies flank the centromere, the chromosome 6 can be represented as a concatenate *ARCRB*, where *A* and *B* refer to the arms of the chromosome, *R* refers to the 300 kb long SD, and *C* refers to the centromere. Canu incorrectly resolved the repeat *R* (resulting in a misassembly *ARB* that erroneously skipped the centromere and collapsed the repeat *R*) while Flye represented the repeat *R* as a single edge in the assembly graph with an elevated depth of coverage (no misassembly).

mosaicFlye recruited 3500 reads (total length 97 Mb) from the repeat *R* and resolved this repeat by leveraging the divergence in the repeat copies (2-3% for most segments) and several insertions of size 0.5-3 kb that distinguish two copies of this repeat (Figure 5). Thus, combining the Flye assembly with the mosaicFlye reconstruction of the repeat *R* resulted in a complete assembly of both arms of chromosome 6, leaving the centromere of this chromosome as the only remaining obstacle to a complete assembly of chromosome 6 in a fully automatic fashion.

**Figure 5.**
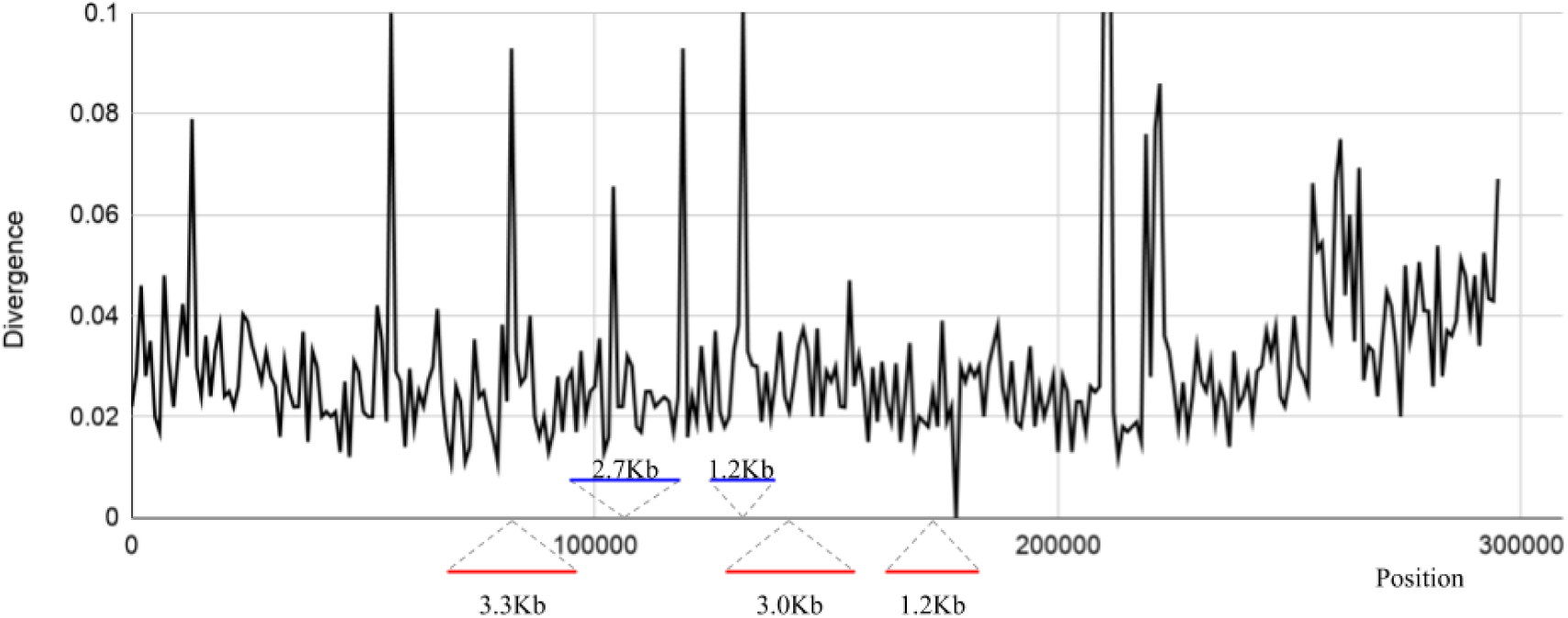
The plot showing the divergence along the two copies of a 300 kb long repeat on chromosome 6. Insertion (deletions) in each copy of the repeat are shown as red (blue) lines at positions corresponding to these indels. Only indels longer than 1 kb are shown.

## Discussion

The ongoing large-scale sequencing projects, such as the Earth Biogenome Project (Lewin et al., 2018) and Telomere-2-Telomere project (Miga et al., 2019), use long-read technologies to generate complete genomes. At the same time, large bacterial sequencing projects (such as NCTC 3000) and metagenomic projects (Bertrand et al., 2019) use long reads to dramatically improve bacterial assemblies. However, since two algorithmic problems (assembling long mosaic repeats and long tandem repeats) remain unsolved, these projects have to complement long read generation by rather expensive complementary technologies such as optical maps, synthetic long reads, and Hi-C reads. Since all these technologies have their inherent error rates (that are still poorly understood), genome assembly is often accompanied by manual analysis that may generate errors. Thus, improving the contiguity of long-read assemblies by resolving mosaic and tandem repeats represent two important goals that will benefit the ongoing genome sequencing projects. We demonstrated that mosaicFlye represents a step in this direction that addresses the key bottleneck in improving the state-of-the-art long-read assemblers. However, it is only the first step since it does not resolve cyclorepeats representing the most complex type of mosaic repeats. Our next goal is to complement the ideas from mosaicFlye with the algorithm from Bzikadze and Pevzner, 2019 that succeeded in resolved some specific cyclorepeats (centromeres) but has not been generalized yet for arbitrary cyclorepeats.

## Appendices

## Fitting and overlap alignments of reads

Given sequences *S_1_* and *S_2_*, we define their maximum-scoring global alignment as *A*=*A*(*S_1_*,*S_2_*). We also define *Query(A) = S_1_* and *Target(A) = S_2_*. The *percent identity PI(A)* is defined as the percentage of matching positions in the alignment *A* among all positions in the alignment. For the sake of simplicity, we ignore chimeric reads that do not align to the genome over their entire length

Given an integer *k* (*k-mer size*), we define *PI_k_(A)* as the minimum of the percent identity among all segments of length at least *k* in the alignment *A* (the default value *k*=1500). Given sequences *S_1_* and *S_2_*, we define their *percent identity* as *PI*(*S_1_*,*S_2_*)=*PI*(*A*(*S_1_*,*S_2_*)), and their *divergence* as *Div(S_1_, S_2_)=100-PI*(*S_1_*,*S_2_*). The concepts *PI_k_*(*S_1_*,*S_2_*) and *Div_k_(S_1_, S_2_)* are defined similarly. Below we use the term “alignment” only for “strong” alignments with sufficiently large percent identity *PI_k_*(*S_1_*,*S_2_*) and ignore all other alignments. Specifically, given a *percent identity threshold PI_min_* (the default value 85%), we say that the sequences *S_1_* and *S_2_ align* if *PI_k_*(*S_1_*,*S_2_*) > *PI_min_*.

Given a read and a genome assembly, various read mapping algorithms (Li, 2018) align this read against all contigs in the assembly. Although reads originating from the unique regions in an assembly are easy to align, it is unclear how to align reads originating from the repeated regions, i.e., to decide which specific copy of a repeat in an assembly a read should be aligned to. To find all such copies, the mapping algorithms usually align such reads to all copies of a repeat (with varying scores) resulting in a set of non-overlapping fitting alignments for each read.

Specifically, given a sequence *S* (a read) and a sequence-set *C* (a set of contigs in an assembly), a “strong” *fitting alignment* of *S* against *C* defines a substring *S’* of one of sequences in *C* that aligns against *S.* i.e., *PI_k_*(*S*,*S’*) > *PI_min_*. Given a sequence *S* and a sequence-set *C*, we describe how mosaicFlye generates the set of all strong non-overlapping fitting alignments *Fitting*(*S, C*). It first finds a highest-scoring fitting alignment of *S* against *C*. If this alignment aligns *S* against a substring *S’* of a sequence *S\** in *C*, we represent *S\** as the concatenate of three sequences *prefix*(*S\**)*, S’*, and *suffix*(*S\**). We further remove *S\** from the sequence-set *C*, substitute it by sequences *prefix*(*S\**) and *suffix*(*S\**), and iteratively repeat the process of finding the strong fitting alignments until it stops, i.e., no strong fitting alignment is found. We refer to the resulting set of fitting alignments as *Fitting(S, C).* In practice, given a read *S* and a sequence-set *C*, we use minimap2 (Li et al., 2018) to generate an approximation of the set *Fitting*(*S, C*). We assume that alignments between reads and genome segments are *transitive* in the following sense: given a read (or a genome segment *S*), if its alignments *A_1_* and *A_2_* belong to *Fitting*(*S, Genome*) then *Target*(*A_1_*) strongly aligns to *Target*(*A_2_*).

Given a read *Read* and a sequence-set *C*, an *overlap alignment* of *Read* against *C* defines a suffix *S’* of one of the sequences in *C* with the maximum scoring global alignment against a prefix *S* of *Read* among all prefixes of all sequences in *C*. We classify an overlap alignment as *strong* if *S* and *S’* have strong alignment and the length of this alignment is at least *k*. If the overlap alignment is strong, we remove the sequence in *C* that has suffix *S’* and iteratively repeat the process of finding the strong overlap alignments until it stops. We refer to the resulting set of overlap alignments as *Overlap(Read, C)* and further combine the sets *Fitting*(*Read, C*) and *Overlap*(*Read, C*) into a single set *Alignments*(*Read,C*). A local alignment between a read *Read* and a contig *c* that aligns a segment *S_1_* of *Read* to a segment *S_2_* of *c* is *correct* if *Origin*(*S_1_*) *= Origin*(*S_2_*).

Appendix “Optimal genome assembly” describes the goal of mosaicFlye with respect to analyzing all fitting and overlap alignments.

## Alignment tournament

Below we assume that each read *Read* was generated from a genome segment *Origin(Read)* and *Fitting*(*Read, Genome*) contains an alignment from *Read* to *Origin*(*Read*) that we refer to as the *correct alignment.* Our goal is to find the correct alignment among all alignments in *Fitting*(*Read, Genome*) between a read *Read* and genome *Genome.* We will first consider the case when the set *Fitting*(*Read, Genome*) includes only two alignments *A_1_* and *A_2_*.

Repeats in a genome accumulate mutations and result in divergent repeat copies, e.g., most segmental duplications in the human genome diverge by more than 1% (Pu et al., 2018). As a result, the correct alignment of a read (to a repeat copy that it originated from) typically has a larger percent identity than the alignment of the same read to an incorrect repeat copy. Moreover, our analysis revealed that, in most cases, the divergence between the repeat copies (defined as *Div*(*Target*(*A_1_*), *Target*(*A_2_*))) is approximately equal to the difference in percent identity *PI*(*A_1_*) *- PI*(*A_2_*) between the correct alignment *A_1_* and an incorrect alignment *A_2_* (Figure A1, left). However, this difference is often low compared to the percent identity between the read and its origin and, in the case of highly similar repeat copies, errors in a read sometimes result in cases when the percent identity of the correct read alignment is lower than the percent identity of an incorrect one (Figure A1).

**Figure A1.**
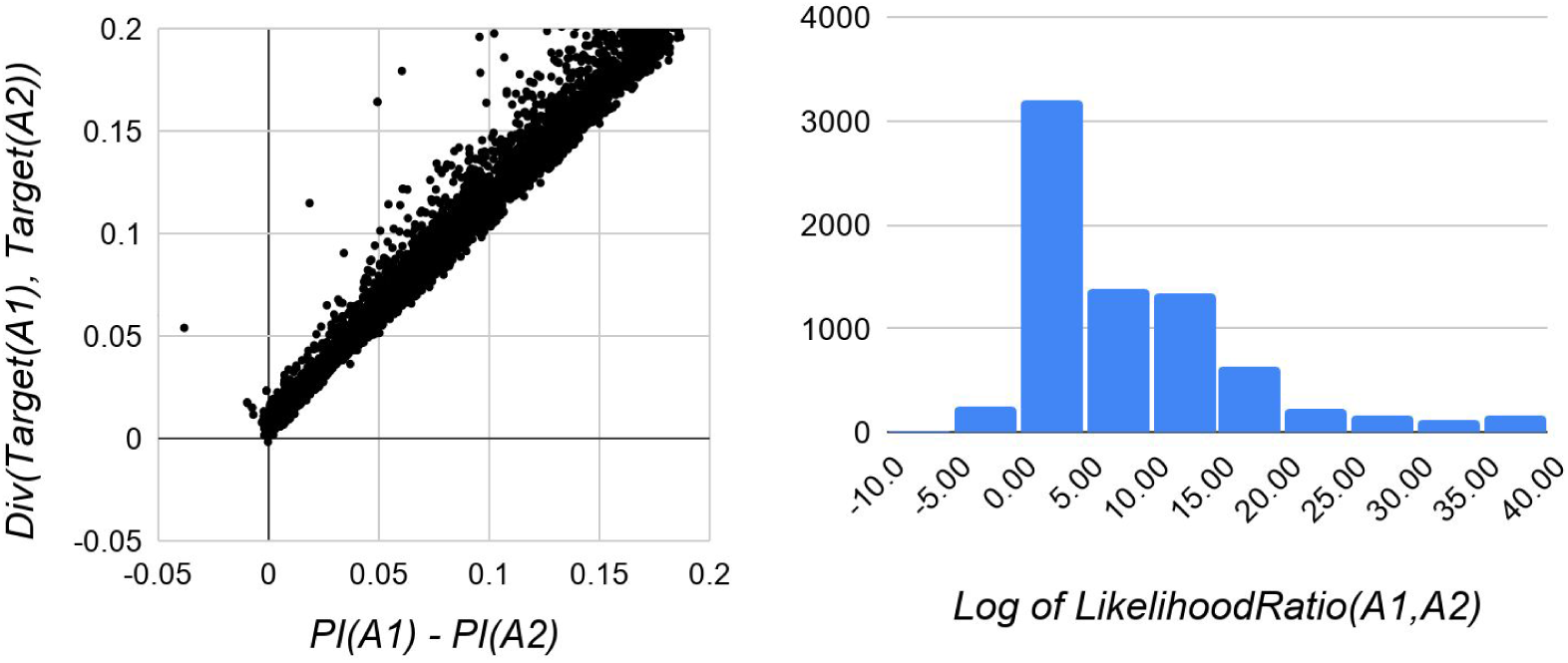
The scatter-plot of the divergence between alignment targets and the difference between percent identities of these alignments (left) and the histogram of the ratio *LikelihoodRatio*(*A_1_, A_2_*) (right). The read alignments were generated for the NCTC dataset 10864.

To avoid selecting false alignments, we use a probabilistic model for comparing alignments described in Lin et al., 2016. Given an alignment *A*, we compute the likelihood that the sequence *Query*(*A*) is generated from a genome segment *Target*(*A*) as a read (for a specific sequencing technology). We further consider the likelihood-ratio *LikelihoodRatio(A_1_*,*A_2_*) to distinguish between two hypotheses: *Origin*(*Read*)*=Target*(*A_1_*) and *Origin*(*Read*)*=Target*(*A_2_)*. If *LikelihoodRatio(A_1_*,*A_2_*) exceeds a threshold *minLikelihoodRatio*, we report *A_1_* as the correct alignment and if falls below *1/minLikelihoodRatio*, we report *A_2_* as the correct alignment. Otherwise, we report no alignment for this read – this usually happens when *Target*(*A_1_*) and *Target*(*A_2_*) have very few diverged positions, making it difficult to infer the correct alignment.

To select the value of *minLikelihoodRatio* we constructed a histogram of the likelihood ratios between the correct alignment *A_1_* and an incorrect alignment *A_2_* for reads from NCTC dataset 10864 (Figure 1A, right). As the histogram illustrates, some incorrect alignments have higher likelihood ratios than the correct alignments. However it hardly ever falls below 10^−5^ which roughly corresponds to the probability of 2-3 mismatches. Based on this observation we chose *minLikelihoodRatio = 10^5^*.

In the case when the set *Fitting(Read, Genome)* contains more than two alignments, mosaicFlye conducts an *alignment tournament* to test each pair of alignments *A_1_* and *A_2_* from *Fitting(Read, Genome)* as described above. An alignment that “won” each pairwise comparison is reported as correct.

We will make the following simplifying assumption about the results of the alignment procedure: if an alignment *A* was selected for a read *Read* then for any segment *S* of *Read* this procedure would either select a reduction of *A* to *S* or return no alignment. We further assume that correct alignments selected by this procedure provide the coverage of the genome with aligned reads that is sufficient for polishing the entire genome and thus *Genome* is a stable point of the alignment-consensus loop described in the next appendix. We will use these assumptions in the mosaicFlye algorithm.

## Optimal genome assembly

Given a read-set *Reads* and a sequence-set *C* we define *fitting* of *Reads* to *C* as a collection of fitting alignments of reads from *Reads* to *C* (at most one fitting alignment for each read). A target-set of a fitting is a collection of targets of all alignments from this fitting. Given an integer *K*, a fitting is called *covering* if each *K*-mer in *C* is contained within at least *minCover* segments from the origin-set (the default value *minCover*=10). Note that the collection of alignments between reads and their origins in *Genome* forms a fitting from *Reads* to *Genome* that we refer to as *origin fitting*. Given a read-set and a *K*-mer from *Genome*, we define its coverage as the number of read origins that contain this *K*-mer. The *K*-mer coverage of a genome is defined as the average coverage of its *K*-mers. mosaicFlye sets the value of *K* in such a way that the *K*-mer coverage is equal to *minCover* (for a typical bacterial genome, *K =* 2500 for a read-set with coverage depth 50x and average read length 6 kb).

A sequence *S* is a *genome candidate* for a read-set *Reads* if there exists a covering fitting of reads from this read-set to *S*. We assume that the origin of each read coincides with one of the target sequences in *Fitting(Read, Genome)* and that the read coverage of the genome is uniform. Thus, *Genome* is also a genome candidate as long as the coverage depth of the read-set exceeds a threshold.

A set of strings is called *free* if no string in the set contains another string as a substring. We define a *genome assembly* as an arbitrary free set of strings and say that an assembly *C_1_ covers* an assembly *C_2_* (written as *C_1_* > *C_2_*) if each sequence from *C_2_* is a substring of a sequence from *C_1_*. An assembly *C* is called *correct* for a read-set *Reads* if for any genome candidate *G* satisfying a condition *AC(G, Reads)*=*G*, we have *C < G*. Strings from a correct assembly are referred to as *contigs*. We will assume that each contig *c* matches to only one segment of the genome and denote this segment as *Origin(c)*. An *optimal assembly* for a given read-set is defined as a correct assembly that covers all other correct assemblies for this read-set. Our goal is to find an optimal assembly.

## Alignment-consensus loop

As described in the main text, correct read alignments are required to polish the draft genome but the polished genome is needed to construct the correct read alignments. To resolve this catch-22, mosaicFlye uses the *alignment-consensus loop* described below. It selects a highest-scoring alignment of each read to the draft assembly and uses the aligned reads for polishing. Given a sequence *S* (a draft genome) and a read-set *Reads* generated from an unknown genome, we refer to a *single* application of this procedure as the *alignment-consensus* (*AC)* step and denote its result as *AC(S,Reads)*.

One can apply the AC step iteratively to a draft error-prone sequence of a genome in a hope that it will converge (Figure A2). We refer to such an iterative application of this procedure as the *alignment-consensus loop* (*ACL).* Although ACL does not necessarily converge, it is not always necessary to find the correct alignment for each read as we only need to align enough reads to provide a sufficient read coverage to polish the genome sequence. As described in Appendix: “Alignment tournament”, the alignment tournament procedure has a low false alignment rate but reports “no alignment” decision in the case of difficult-to-align reads.

**Figure A2.**
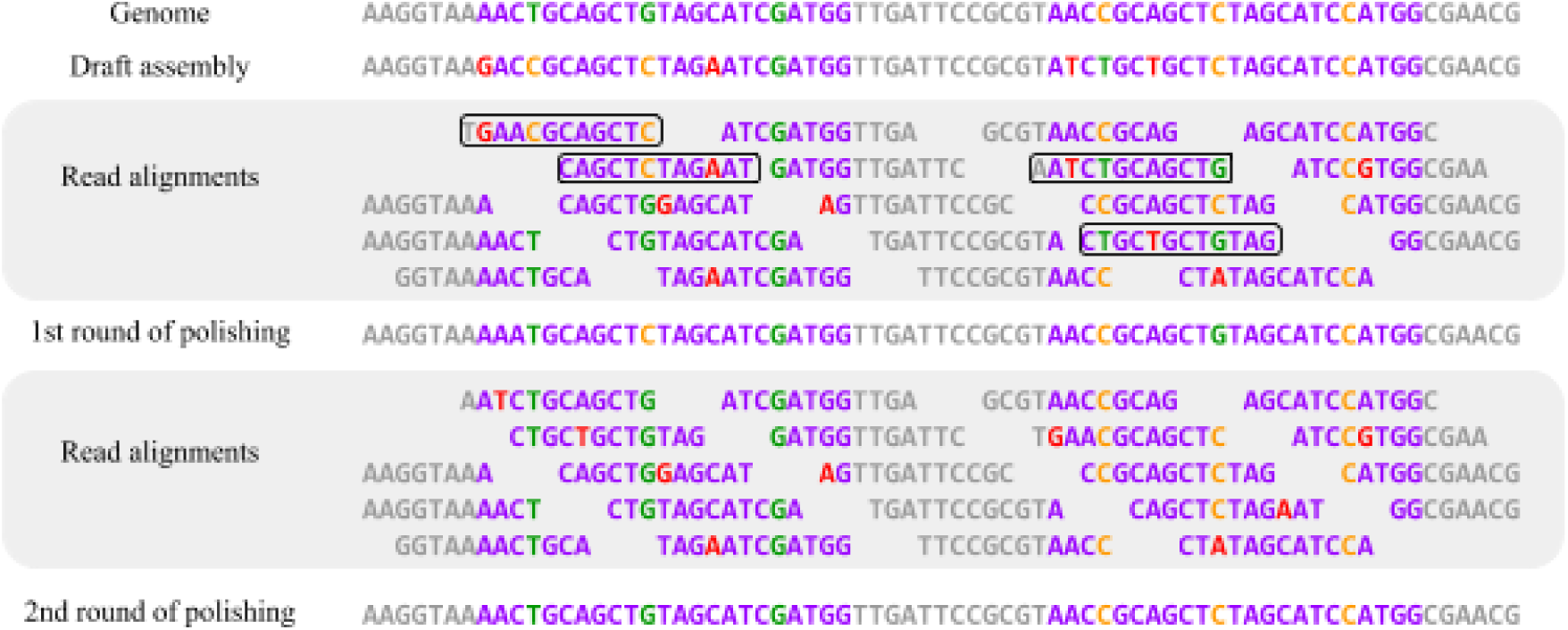
The alignment-consensus loop. Single nucleotide variations between two copies of the “purple” repeat are shown in green (in the first copy) and in orange (in the second copy). Reads originating from the first (second) copy of this repeat “inherit” green and orange variations. Errors in the draft assembly and reads are shown in red. Four reads that have incorrect alignments to the draft assembly at the first round of polishing are shown in solid boxes. The same four reads that have correct alignments to the draft assembly at the second round of polishing are shown in solid boxes.

## Expanding alignment-consensus loop

Instead of considering a complete genome, we will now consider a genome assembly and apply the ACL for genome assembly rather than for polishing as before.

We define the *multiplicity* of a string *S* in a genome *Genome* as the size of *Fitting(S, Genome)* and refer to sequences of multiplicity one as *unique*. The Flye assembler uses reads to construct the *repeat graph* of the genome and further uses *bridging* reads to resolve the *repeat edges* in this graph (Kolmogorov et al., 2019). mosaicFlye starts from the Flye repeat graph (before the repeat resolution step that uses the bridging reads) and selects sequences of all unique edges in this graph as the contigs in the initial mosaicFlye assembly *C_0_* (shown in grey in Figure A3). Since Flye collapses all repeats of length *k* and longer, the unique edges in the Flye repeat graph contain only unique *k*-mers, i.e., *k*-mers that do not align to other regions in *C_0_*.

**Figure A3.**
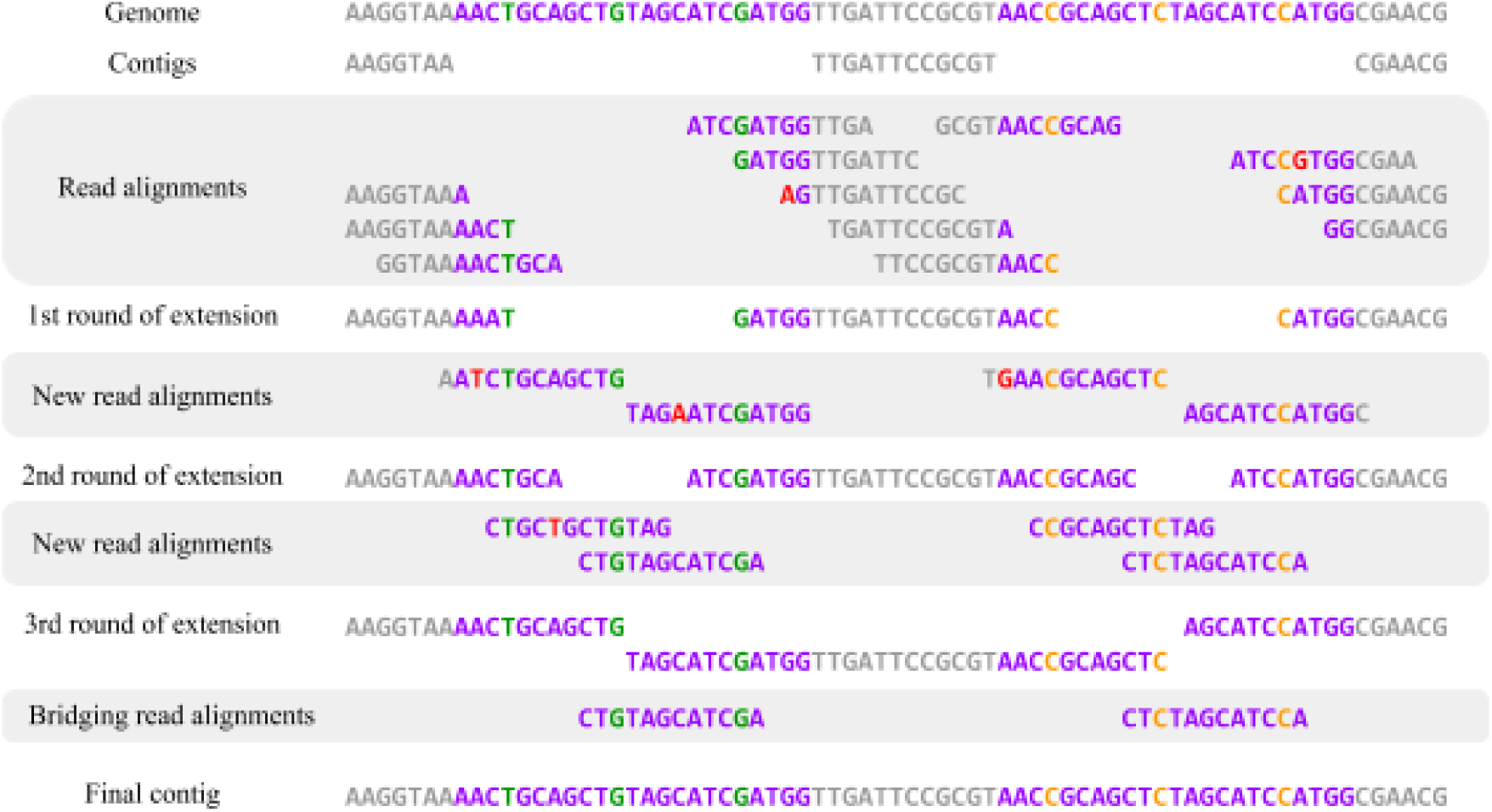
The expanded alignment-consensus loop reconstructs two instances of a repeat.

mosaicFlye uses parameters *k* and *K* (*k < K*) and assumes that all reads have length at least *k*. Since all *k*-mers in assembly *C_0_* are unique, all alignments of reads to sequences from *C_0_* are correct. Thus, we can assume that sequences in *C_0_* are polished and *C_0_* is a correct assembly. mosaicFlye constructs a series of correct assemblies *C_0_ < C_1_ < C_2_ … < C_m_ = C_m+1_* until this process converges, i.e., until *C_m_ = C_m+1_* (*C_m_* is reported as an approximation of an optimal assembly). At each step, mosaicFlye constructs correct alignments of reads to sequences in an assembly (*contigs*) and uses these reads to construct even longer contigs as illustrated in Figure A3. We refer to this procedure as the *expanding alignment-consensus loop* (*EACL*).

We start by explaining how EACL constructs *C_1_* from *C_0_*. Given a contig *c* from *C_0_* and a read *Read* mapped to this contig, all alignments from *Overlap(Read, c)* are correct since all *k*-mers from *C_0_* are unique. Since any *K*-mer in the genome is covered by at least *minCover* reads then at least *minCover* reads that cover the last *k*-mer of *c* also cover the next *K – k* nucleotides in *Genome* after *Origin(c)*. Thus, the contig *c* in *C_0_* can be prolonged by at least *K – k* nucleotides using one these reads. Since the resulting expanded contig has sufficient coverage (for at least *K – k* nucleotides), it can be polished over its entire length resulting in longer polished contigs as compared to *c*. Applying this procedure to all contigs in *C_0_* results in the set of elongated contigs *C_1_*. The consequent assemblies *C_2_, C_3_,…* are constructed in a similar way by selecting the correct alignment for reads that overlap with contigs and using new reads to prolong these contigs.

## Expanding alignment-consensus loop for transforming the repeat graph into the de Bruijn graph

The alignment-consensus loop can be applied to a draft assembly resulting from a traversal of the repeat graph. If we knew which reads originated from each copy of a repeat edge (for each repeat edge in the repeat graph), we would be able to accurately polish sequences of each copy of each repeat and thus reconstruct the de Bruijn graph as described in the previous sub-section. After constructing the de Bruijn graph, one can find the top-scoring alignment of reads to paths in the de Bruijn graph using an approach similar to the hybridSPAdes algorithm (Antipov et al., 2016). Similarly to the top-scoring alignment of a read against the genome, the top-scoring alignment of a read against the de Bruijn graph is likely to be correct (it is still impossible to find the correct alignment for reads that align to several identical or nearly identical copies of a repeat).

mosaicFlye applies the EACL to gradually transform the repeat graph into the de Bruijn graph and thus to resolve repeats by revealing unique edges in the de Bruijn graph that were collapsed by the Flye algorithm for the repeat graph construction. We classify an edge of a graph as *resolved* if its sequence is a substring of a genome and as *unique* if it is resolved and occurs only once in the genome. In the de Bruijn graph all edges are resolved. In the repeat graph, unique edges are resolved and unique (since they can be polished using correct alignments) but repeat edges are not resolved. mosaicFlye constructs a sequence of graphs *G_0_ = RG(Reads, k), G_1_, G_2_,…* where each graph contains all resolved edges of the previous graph as substrings of its resolved edges. At each step, mosaicFlye constructs correct alignments of reads on resolved edges and uses these reads to prolong the perfectly polished sequence and create more resolved edges (Figure 3 in the main text).

## The challenge of aligning reads to the growing assembly

The EACL approach, while conceptually simple, faces the challenge of selecting the correct alignment of reads to the growing contigs. We already discussed this problem in two settings: selecting correct alignment of a read to a genome and to an assembly *C_0_*. We will combine techniques used in these two cases to solve a more difficult problem of aligning a read to an assembly *C_i_* for *i=1, …, m*.

Although aligning a read to an assembly *C_0_* is an easy task (since all *k*-mers in *C_0_* are unique), it is not clear how to align a read to an assembly *C_1_* since some *k*-mers in *C_1_* are non-unique. Indeed, since a read may overlap with several contigs in *C_1_*, the choice of the correct overlap alignment becomes non-trivial. Also, it is not clear how to generalize the previously described approach for aligning a read against a (complete genome) to aligning it against an (incomplete) assembly.

To address this challenge, we will generalize the concept of a unique *k*-mer. Let *C* be a correct assembly of a genome *Genome*. We say that a sequence *S* is *resolved* with respect to the assembly *C* if all genomic segments it aligns to are contained within contigs in *C*, i.e., each segment from *Target(Alignments(S, Genome))* is a segment of *Origin(C)*. Another equivalent definition of a resolved sequence is as follows: sequence *S* is resolved if |*Fitting(S,C)*| *=* |*Fitting(S,Genome)*|. Figure A4 shows an example of a resolved segment *S_1_* (both genomic segments *S_1_* aligns to are contained within contigs in *C*) and an unresolved segment *S_2_* (one of two genomic segments it aligns to is not contained within a contig in *C)*.

**Figure A4.**
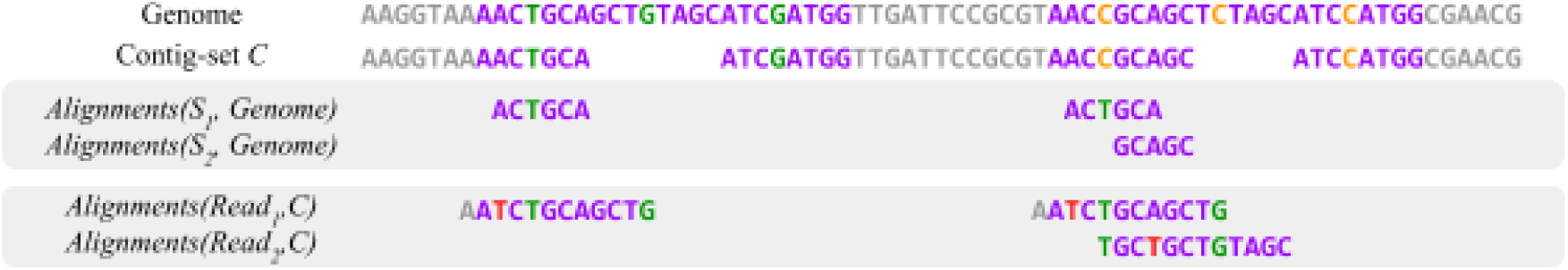
Resolved segments and correct read alignments. A segment *S_1_* is resolved with respect to assembly *C* because all genomic segments it aligns to are contained within contigs in *C*. In contrast, a segment *S_2_* is not resolved because one of two genomic segments it aligns to is not contained within a contig in *C*. A segment of a read *Read_1_* aligns to a resolved segment *S_1_* and thus one of its alignments to contigs from assembly *C* is correct. In contrast, since *Read_2_* does not align to a resolved segment, its alignment to assembly *C* is incorrect.

If a read *Read* aligns to a resolved segment *S* in an assembly *C*, then *Origin(Read)* is a segment of *Origin(C)* since, by the alignment transitivity condition, *Origin(Read)* aligns to a segment of *Origin(C)* and *S* is resolved. Moreover, even if only a segment *S* of a read aligns to a resolved segment of *C*, *Origin(S)* is a segment of *Origin(C)* and *Read* has either a correct fitting alignment or a correct overlap alignment with *C*. Figure A4 shows an example of a read *Read_1_* that contains a segment that aligns to a resolved segment *S_1_* in assembly *C* and thus has a correct overlap with *C*. In contrast, a read *Read_2_* does not have a correct overlap with contigs and none of its segments aligns to any resolved segment. Thus, finding the highest-scoring alignment among all alignments in *Fitting(S, C)* would report the correct alignment of *S* to *C* that can be expanded to correct overlap or fitting alignment of *Read* to *C*.

Unfortunately, it is not possible to directly check if a read or its segment is resolved when the genome is unknown. Below we show how to find resolved segments of contigs (without knowing the genome sequence) by keeping track of resolved sequences of length *k*.

## Expanding the set of resolved *k*-mers by traversing mosaic repeats

Flye glues all copies of *long repeats* (i.e., repeats of length at least *k*) into a single edge in the repeat graph, and further classifies all edges of the repeat graph into unique (that represent unglued genomic regions) and repeat edges. Removing unique edges from the repeat graph reveals the connected components formed by the repeat edges that we refer to as *mosaic repeats*. Since the sequence of a repeat edge represents a consensus sequence of all repeat copies, it typically deviates from sequences of these copies. We will first describe a simple case of mosaic repeats that do not have cycles (*acyclic mosaic repeats*) and later consider mosaic repeats that have cycles (referred to as *cyclorepeats*)

Flye reconstructs the accurate sequences of unique edges that we use as the initial contigs in the EACL approach. Each *k*-mer from a unique edge is unique and thus has only one alignment to the genome. Therefore, since each such *k*-mer occurs in one of the contigs, all such *k*-mers are resolved. We will initialize the set of *resolved k-mers R_0_(k)* as the set of all *k*-mers occurring in unique edges. Since reads from long repeats do not contain unique *k*-mers, we will iteratively expand the collection *R_0_(k)* with each EACL iteration in an attempt to find as many resolved *k*-mers as possible, thus creating a series of *k*-mer sets *R_0_(k) ⊂ R_1_(k) ⊂ R_2_(k) ⊂… ⊂ R_m_(k)*, where *R_i_(k)* consists of resolved *k*-mers in an assembly *C_i_* for *i* = 0,…,*m*. We will find these *k*-mers as segments of the newly constructed contigs.

We assume that *Genome* has a strong alignment to a path (referred to as a *genome traversal*) in the repeat graph and refer to the number of times this traversal passes an edge *Edge* in the repeat graph as *Multiplicity(Edge)*. Since all repeats of length at least *k* are glued in the repeat graph, the multiplicity of a *k-*mer from an edge *Edge* is equal to *Multiplicity(Edge)*. Given an acyclic mosaic repeat, one can infer the multiplicity of each edge in this graph (Kolmogorov et al., 2019a) and thus the multiplicity of each *k*-mer. Thus, a *k*-mer *S* occurring on the edge *Edge* is resolved in an assembly *C* iff *Fitting(S, C) = Multiplicity(Edge)*.

Figure A5 shows an example of an iterative expansion of contigs and a parallel expansion of the sets *R_i_(k)* (shown by dashed lines) and assemblies *C_i_* (shown in purple). The initial contigs (top left) correspond to unique (grey) edges and the initial set *R_0_(k)* corresponds to all *k*-mers in the initial contigs. In the first step, contigs are extended inside the repeat based on reads that share unique *k*-mers with unique edges (top right). The structure of the repeat graph reveals that the multiplicities of edges A and B are 2 and 3, respectively. Thus, all *k*-mers from the edge A, that are covered by both contigs that extended inside A, are resolved. However, no *k*-mers from edge B can be marked as resolved since they are covered by at most one contig. Thus, at this point, we can only align reads to two of the three contig and continue extending these contigs inside the edge A. As the two contigs are extended into the repeat edge A, the collection of *k*-mers covered by these contigs also expands allowing to align more reads and prolong contigs further until the entire edge A is resolved (middle left). After all three contigs prolong into the edge B, we can mark some of the *k*-mers from B as resolved (middle right), eventually restore all copies of the repeat edge B, and finally find out how to connect these contigs with outgoing edges (bottom).

**Figure A5.**
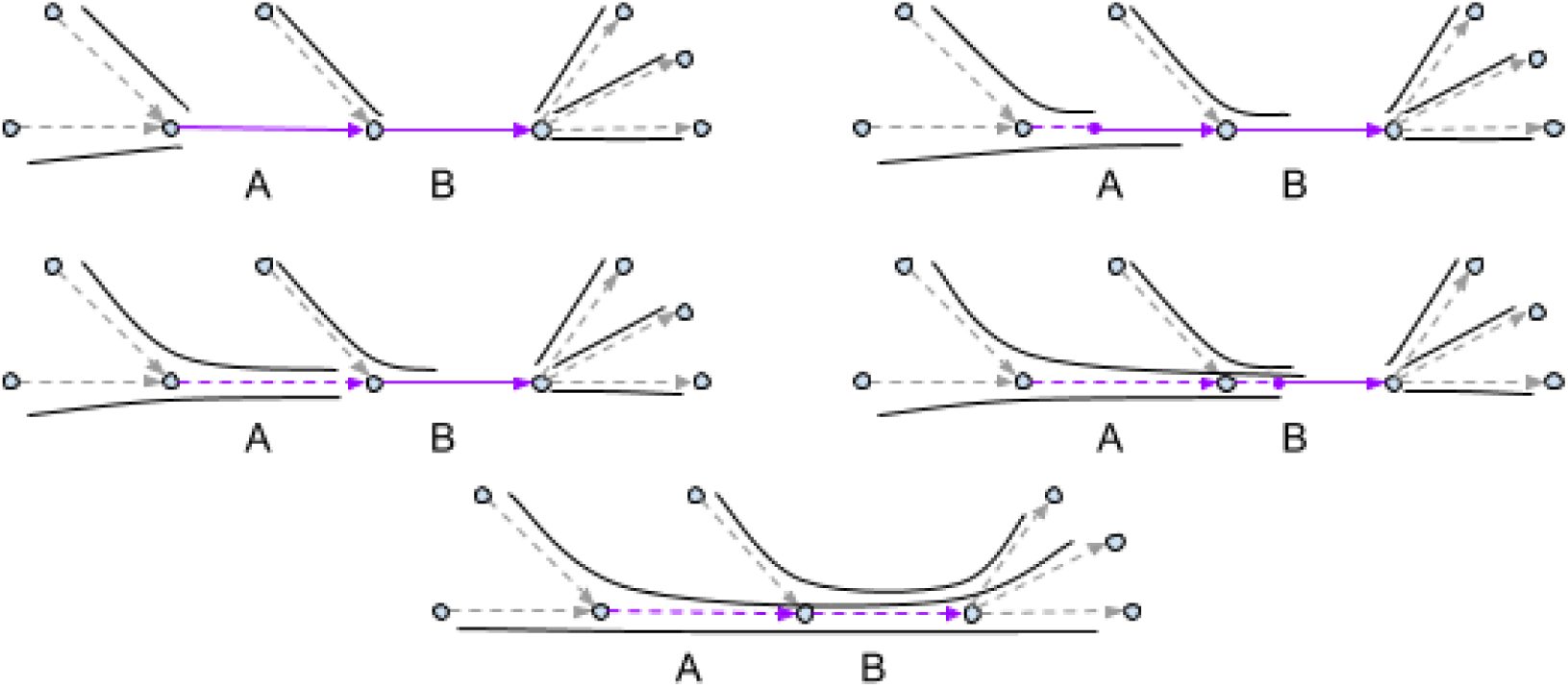
An iterative expansion of contigs with a parallel expansion of the *k*-mer sets *R_i_(k)*. Sets *R_i_(k)* are formed by all *k*-mers from the dashed edges. All edges in the graph are unique except for edge A with multiplicity 2 and edge B with multiplicity 3.

## Corrupted mosaic repeats

The described algorithm works well in the case when Flye accurately represents a mosaic repeat but faces difficulties in the case of *corrupted* mosaic repeats that inaccurately represent various copies of a mosaic repeat. Moreover, it requires accurate estimates of multiplicities of edges in a mosaic repeat that are often difficult to obtain even in the case of non-corrupted mosaic repeats (Kolmogorov et al., 2019).

To deal with errors in long reads, Flye aggressively collapses bulges (pairs of edges connecting the same vertices) and contracts short edges in the repeat graph. As a result, the genome sequence may not have a strong alignment to a genome traversal resulting from the Flye repeat graph. Moreover, after bulge collapsing, some *k*-mers from repeat copies may become so diverged from the consensus that the expansion of the sets *R_i_(k)* may fail since reads do not align to any consensus *k*-mers. Below we describe a method to expand the *k*-mer-set set without relying on the repeat graph in such a way that *R_i_(k)* consists of *k*-mers in expanding contigs rather than from the *k*-mers from the repeat edges in the Flye assembly that represents a consensus of multiple repeat copies.

Note that if a segment *S* is resolved in an assembly *C* then a segment *S’* from this assembly that contains *S* as a substring is often also resolved. Indeed, since all alignments of *S* to the genome are covered by *Origin(C)* then all alignments of *S’* are at least partially covered by *Origin(C)*. Using this observation, we will show how to (i) construct a set of longer resolved *K*-mers than the original set of resolved *k*-mers, and (ii) find additional resolved *k*-mers using the set of resolved *K*-mers.

Specifically, given a set *R(k)* of resolved *k*-mers in an assembly, we will construct a set of resolved *K*-mers *R(K)* in a new assembly, where *k*-prefixes of all *K*-mers from *R(K)* belongs to *R(k)*. Afterward, we will find *K*-mers from *R(K)* whose *k-*suffixes represent resolved *k*-mers that do not belong to *R(k)*. Such resolved *K*-mers connect a resolved *k-*prefix with a resolved *k*-suffix, not unlike how a *(k+*1)-mer connects its *k-*prefix and *k*-suffix to form an edge in the conventional de Bruijn graph. Finally, we will iteratively expand the set *R(k)* by adding all such *k*-suffixes.

## From a resolved prefix *k*-mer to a resolved *K*-mer

We say that a *K*-mer *S extends* a *k*-mer *s* if *s* is a prefix of *S*. Consider a *K*-mer *S* in an assembly *C* that extends a resolved *k*-mer *s*. We will show that if *Overlap(S, C)* is empty then *S* is resolved, i.e., any “strong” alignment of *S* to contigs is a fitting alignment rather than an alignment of a prefix of *S* to a suffix of a contig from *C*.

Indeed, any alignment *A* of *S* to *Genome* also aligns *s* to *Genome* (Figure A6). Since *s* is resolved, there is an alignment *A’* in *Alignments(s, C)* that aligns the *k-*mer *s* to a contig *c* such that a segment *Origin(Target(A’))* is a prefix of segment *Target(A)*. Since *Origin(c)* contains *Origin(Target(A’))*, either *Origin(c)* contains *Target(A)* or *Target(A)* overlaps *Origin(c)*. In the latter case prefix of *S* aligns to a suffix of *c*, a contradiction with the initial assumption. Thus *Origin(c)* contains *Target(A)*. Consequently, every alignment of *S* to *Genome* is covered by one of the alignments of *S* to contigs and *S* is resolved.

**Figure A6.**
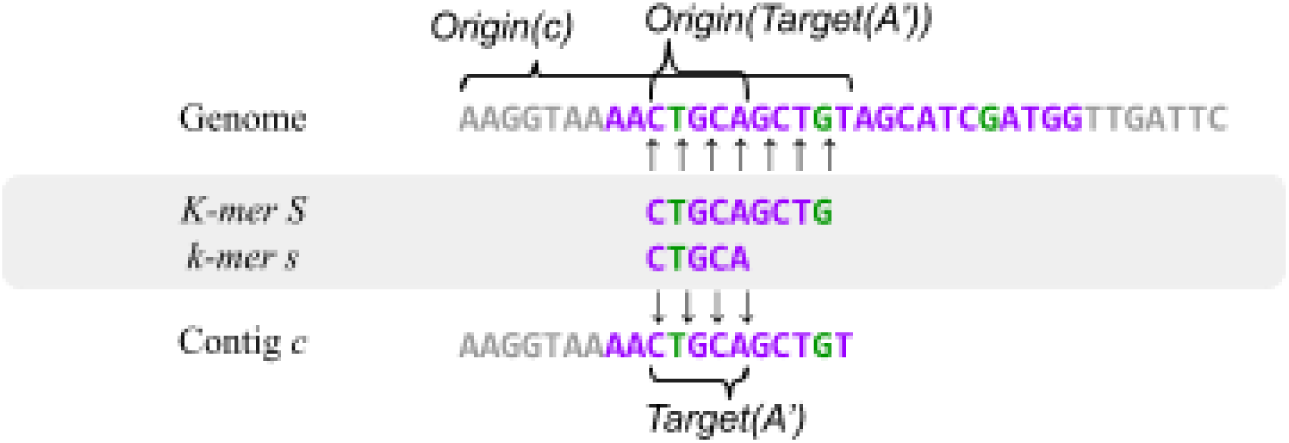
From a resolved *k*-prefix to a resolved *K*-mer. A resolved *k*-mer *s* is prolonged to a resolved *K*-mer *S* using alignments of *s* to contigs.

## From a resolved *K*-mer to a resolved suffix *k*-mer

Let *RightExpansion(s, K, C)* (*LeftExpansion(s, K, C))* be the set of all *K*-mers from an assembly *C* which have a *k*-prefix (*k*-suffix) that aligns to a *k*-mer *s* from contigs. We classify a resolved *k*-mer *s* in an assembly *C* as *strongly resolved* if each *K*-mer *S* from *RightExpansion(s, K, C)* satisfies the condition that *Overlap(S, C)* is empty. We have shown that if a *k*-mer *s* is strongly resolved in an assembly *C*, then all *K*-mers from *RightExpansion(s, K, C)* are resolved.

The statement above describes how strongly resolved *k*-mers allow one to identify resolved *K*-mers. We will now formulate an “opposite” statement that describes how resolved *K*-mers allow one to find resolved *k*-mers, at least in the case the genome *Genome* is known. Specifically, if *LeftExpansion(s, K, Genome)* contains only resolved *K*-mers with respect to an assembly *C* then *s* is resolved. Indeed each segment from *Target(Alignments(s, Genome))* is covered by the origin of one of the resolved *K-*mers from *LeftExpansion(s, K, Genome)* and thus it is also covered by *Origin(C)*.

Unfortunately, it is unclear how to check the condition “if *LeftExpansion(s, K, Genome)* contains only resolved *K*-mers with respect to an assembly *C”* when *Genome* unknown. Indeed, we only know which *K*-mers from *C* are resolved but do not know which *K*-mers from *Genome* are resolved. To address this complication, we describe a criterion that uses read alignments to check whether *LeftExpansion(s, K, C) = LeftExpansion(s, K, Genome)* in the case when *LeftExpansion(s, K, C)* contains only resolved *K*-mers.

Let *Reads(s, K)* be the set of all reads whose origin covers one of the *K*-mers from *LeftExpansion(s, K, Genome)*. These reads can be detected even without *Genome* as all reads that have a fitting alignment with *s* that starts at or after the position *K – k* in the read (Figure A7). Since all *K*-mers in the genome are covered by at least *minCover* reads, segments from *Origin(Reads(s, K))* cover each *K*-mer from *LeftExpansion(s, K, Genome)* at least *minCover* times.

**Figure A7.**
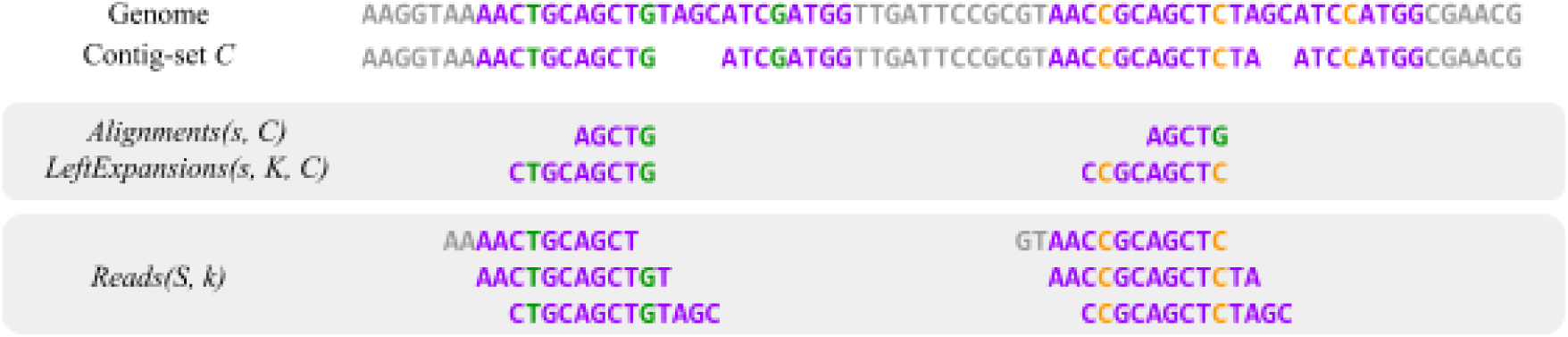
From a resolved *K*-mer to a resolved *k*-mer. *LeftExtensions(s, K, C)* for the *k*-mer *s=AGCG* consists of two resolved *K*-mers: CTGCAGCG and CCGCAGCTC. *Reads(S, k)* consists of 6 reads each of which aligns to one of the left expansions of *s*. In this case, we can conclude that *k*-mer *s* is resolved.

Consider a *K*-mer *S* in *LeftExpansion(s, K, Genome)* that does not belong to *LeftExpansion(s, K, C)* and a *Read* such that *Origin(Read)* contains *S*. We will show that in this case *Read* does not have a fitting or an overlap alignment with *C*. If *Read* aligns or overlaps with a contig *c* from *C* then, by the transitivity condition, a *K*-mer *S’* from *c* aligns to *S*. This alignment can be reduced to a strong alignment of the suffix of *S’* of length *k* to the suffix of *S* that aligns to *s*, implying that the suffix of *S’* aligns to *s* and *S’* belongs to *LeftExpansion(s, K, C)*. We assumed that all *K*-mers from *LeftExpansion(s, K, C)* are resolved, implying that *S’* is also resolved. Since *S’* aligns to *S*, *S* is covered by contigs, a contradiction to the assumption that *S* does not belong to *LeftExpansion(s, K, C)*. Thus, *Read* can not have a fitting or an overlap alignment with *C*. This conclusion leads to the following test for deciding whether *s* is resolved: if all *K*-mers from *LeftExpansion(s, K, C)* are resolved and *Alignments(Read, C)* is empty for each read *Read* in *Reads(s, K)*, then *s* is resolved.

## Merging contigs and bridging repeats

As we extend contigs using the EACL, origins of previously non-overlapping contigs may eventually overlap, creating an opportunity to merge these contigs in an assembly. We say that a read *Read connects* contigs *c_1_* and *c_2_* if (i) there exists an overlap alignment *A*(*c_1_,Read*) of contig *c_1_* with *Read*, (ii) there exists an overlap alignment *A*(*Read,c_2_*) of *Read* with contig *c_2_*, and (iii) the corresponding overlap alignments satisfies the condition that *Query(A*(*c_1_,Read*)) overlaps with *Query(A*(*Read,c_2_*)). We say that a read *Read scaffolds* contigs *c_1_* and *c_2_* if conditions (i) and (ii) hold but the condition (iii) does not hold.

Since a connecting read is an indication that the origins of contigs *c_1_* and *c_2_* overlap (note that EACL constructs reliable alignments), mosaicFlye merges these contigs into a single one. A scaffolding read provides evidence that two contigs are separated by a gap sequence in the genome even though it does not provide sufficient information to accurately infer this sequence in the case there are few scaffolding reads between a pair of contigs. Nevertheless, mosaicFlye utilizes information about scaffolding reads by performing an additional *scaffolding* step after the EACL converges. The (potentially error-prone) gap sequence between the scaffolded contigs is computed as the consensus derived from gap sequences of all scaffolding reads for these contigs.

## Dataset details

Appendix: Dataset details.
| Dataset name | Coverage | Median read length | Genome size (Mb) | #Contigs in Flye assembly |
| --- | --- | --- | --- | --- |
| 4450 | 131 | 3056 | 5.5 | 29 |
| 7921 | 102 | 3470 | 5.3 | 5 |
| 8294 | 224 | 3657 | 2.9 | 4 |
| 8333 | 67 | 3142 | 5.6 | 8 |
| 8359 | 273 | 2042 | 3.0 | 8 |
| 8603 | 111 | 3406 | 5.1 | 4 |
| 8684 | 83 | 2431 | 4.8 | 10 |
| 9002 | 135 | 3356 | 5.0 | 3 |
| 9006 | 86 | 4384 | 5.1 | 3 |
| 9007 | 105 | 4653 | 5.3 | 4 |
| 9012 | 78 | 4428 | 5.1 | 4 |
| 9016 | 73 | 4970 | 5.0 | 8 |
| 9024 | 66 | 3557 | 5.3 | 4 |
| 9028 | 102 | 2203 | 5.1 | 10 |
| 9657 | 61 | 3044 | 5.1 | 7 |
| 10864 | 88 | 3452 | 5.2 | 9 |
| 11658 | 190 | 1830 | 2.5 | 6 |
| 11692 | 54 | 2640 | 5.3 | 4 |
| 11962 | 221 | 3102 | 2.9 | 3 |
| 12158 | 34 | 3628 | 5.9 | 5 |

## Time complexity

Each iteration of the contig expansion requires *O(m^2^RC)* time where *R* is the maximal read length, *m* is the multiplicity of a repeat and *C* is genome coverage. If the iteration results in very small extension of length < *d* (we use d = 500) then we stop expansion of contigs thus the number of iterations does not exceed the total length of all repeat copies (*L*) divided by *d* resulting in overall running time *O(m^2^RCL/d)*. However, this theoretical time complexity is further optimized in the code. As a result resolution of 300Kb long repeat with coverage 115 by reads with N50 70Kb in the human genome takes 30 hours.

